# Amyloid plaques drive state-dependent long-range circuit reorganization in the hippocampus

**DOI:** 10.64898/2026.04.22.720180

**Authors:** Z Zhao, JJ Locascio, H Li, N Gowravaram, RJ Green, K Kastanenka, B Bacskai, BT Hyman, SN Gomperts

## Abstract

Amyloid plaques are a pathological hallmark of Alzheimer’s disease, but how they drive widespread neuronal dysfunction remains unclear. While studies in anesthetized animals show that plaques drive local hyperactivity^1,2^, it is unknown how this pathology shapes functional hippocampal maps in freely behaving animals. We combined chronic 1-photon calcium imaging, local field potential recordings, and post hoc 2-photon plaque imaging in freely behaving APP/PS1 mice across behavior and sleep to correlate real-time hippocampal activity and place coding with precise plaque topography. Here we show that plaques exert nonlocal, long-range effects on hippocampal activity that depend on plaque size, laminar position, and the animal’s behavioral state. Place cells, which encode spatial position and are normally uniformly distributed, are preferentially enriched near plaques, revealing an aberrant reorganization of plaque-adjacent neurons into the hippocampal map of space. In longitudinal experiments, pre-existing place cell locations do not predict future plaque sites, whereas hyperactivity during slow-wave sleep weakly predicts future amyloid deposition. These findings identify a mechanism by which amyloid pathology reorganizes brain circuits, degrading the functional architecture of the hippocampus and contributing to widespread dysfunction and cognitive impairment in Alzheimer’s disease.

---

During exploratory behavior, many hippocampal neurons form stable spatial representations known as place fields. Notably, spatial representations deteriorate in parallel with spatial memory as plaques accumulate in murine amyloid-β_42_ (Aβ) models of Alzheimer’s disease^3–5^, mimicking early navigational deficits in patients (e.g., ^6^). Aberrantly hyperactive and silent neuronal subpopulations in hippocampus and cortex of murine Alzheimer’s disease models suggest that disrupted excitatory-inhibitory balance underlies Aβ-associated circuit dysregulation^1,2,7^. Fibrillar and oligomeric Aβ species are both neurotoxic and synaptotoxic ^8–14^ and are well-positioned to disrupt circuit function.

In anesthetized Alzheimer’s disease model mice, aberrantly hyperactive cells have been found to cluster exclusively near amyloid plaques, while silent cells have been found to be uniformly distributed^1,2^. However, behavioral state critically influences hippocampal activity^15,16^, and anesthetics have complex effects on hemodynamics, neurophysiology, and calcium homeostasis^17–19^. Therefore, whether plaque-associated hyperactivity occurs during normal behaviors and across the sleep-wake cycle, and how it impacts place coding, remains unknown. Furthermore, the impact of aberrant clustering of hyperactive cells proximal to plaques on place coding has not been established; this clustering and hyperactivity is likely to undermine place coding and degrade pattern separation needed for accurate storage of unique experiences.

To address these gaps, we quantified CA1 calcium dynamics and concomitant electrophysiological activity in freely-behaving APP/PS1 mice, and then co-registered these data to high-resolution plaque topography. We show that plaques induce nonlocal, long-range, state-dependent effects on neuronal activity and place coding that depend on plaque size and laminar location within CA1. These findings provide a new framework in which amyloid plaques act as foci of network-level circuit reorganization, rather than as purely focal lesions.

## Behavioral state transforms the spatial influence of amyloid plaques on hippocampal circuits

To investigate how amyloid plaques impact hippocampal function across behavioral states, we acquired simultaneous chronic GCaMP6f calcium imaging and local field potential (LFP) recordings in the CA1 region of freely behaving APP/PS1 mice across the sleep-wake cycle, followed by high-resolution plaque imaging (**Fig. 1a,b** and **Supplementary Fig. 1**; see **Methods**). We stratified neurons into three activity classes, high activity, moderately active, and quiescent cells, based on state-specific calcium event rates (**Fig. 1b,c**).

**Figure 1.**
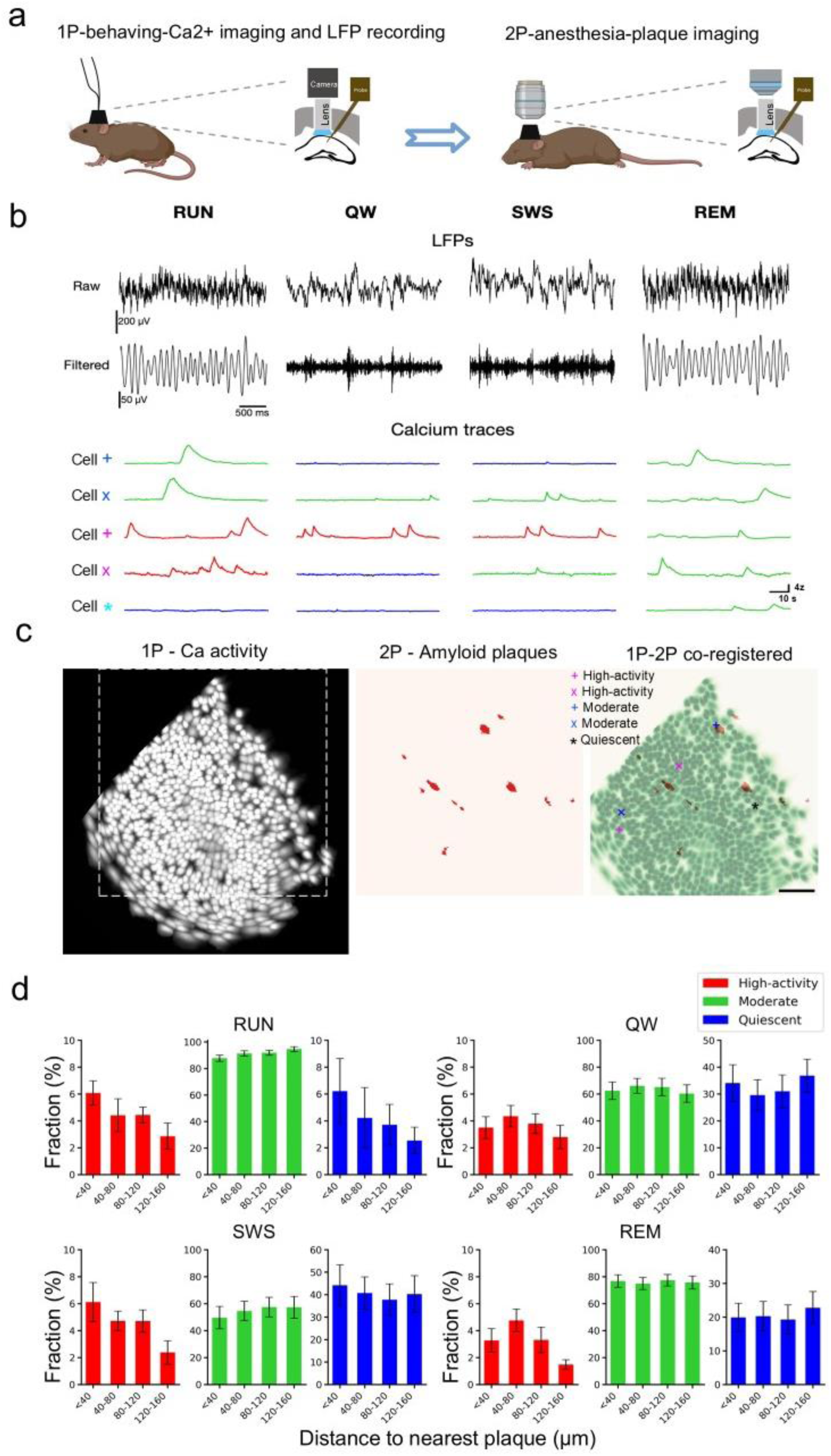
Plaque topography modulates hippocampal ensemble activity across behavioral states. **a**, Experimental workflow: simultaneous 1-photon (1P) miniscope hippocampal dynamic calcium imaging and local field potential (LFP) recording in CA1 of freely behaving APP/PS1 mice, followed by *in vivo* 2-photon (2P) imaging of amyloid plaques. **b**, Representative LFPs (top) and corresponding calcium transients (bottom) during exploratory behavior (RUN), quiet wakefulness (QW), slow-wave sleep (SWS), and rapid eye movement sleep (REM) sleep. Neurons are stratified by activity: high-activity (red), moderate (green), and quiescent (blue). **c**, 1P-imaged GCaMP6f somatic activity (left, white) co-registered with 2P-imaged Methoxy-X04 labeled plaques (middle, red). Right: Overlay of somas (green) and plaques (red) with neurons from (**b**) indicated. **d**, Proportion of quiescent (blue), moderate (green), and high-activity (red) neurons as a function of distance from the nearest plaque (n=7 mice) across behavioral states. No significant correlations were observed between distance and the proportion of cells in any activity stratum (Spearman correlation, Bonferroni-adjusted). Data are mean ± s.e.m. (n=7 mice, 3762 cells).

In contrast to previous observations made in anesthetized Alzheimer’s mice^1,2^, high-activity cells were not restricted to the immediate vicinity of amyloid plaques, but rather were detected at all distances from the nearest plaque across all behavioral states (**Fig. 1d and Supplementary Fig. 2**). Although the proportion of high-activity cells visually decreased with distance from the nearest plaque during running and slow-wave sleep, Spearman correlations between distance and the fraction of cells in each activity class were not significant in any state after Bonferroni correction (**Fig. 1d**). By contrast, CA1 activity profiles varied significantly with behavioral state (**Fig. 2a**), indicating that behavioral state, rather than local plaque pathology, primarily determines the distribution of activity strata.

**Figure 2.**
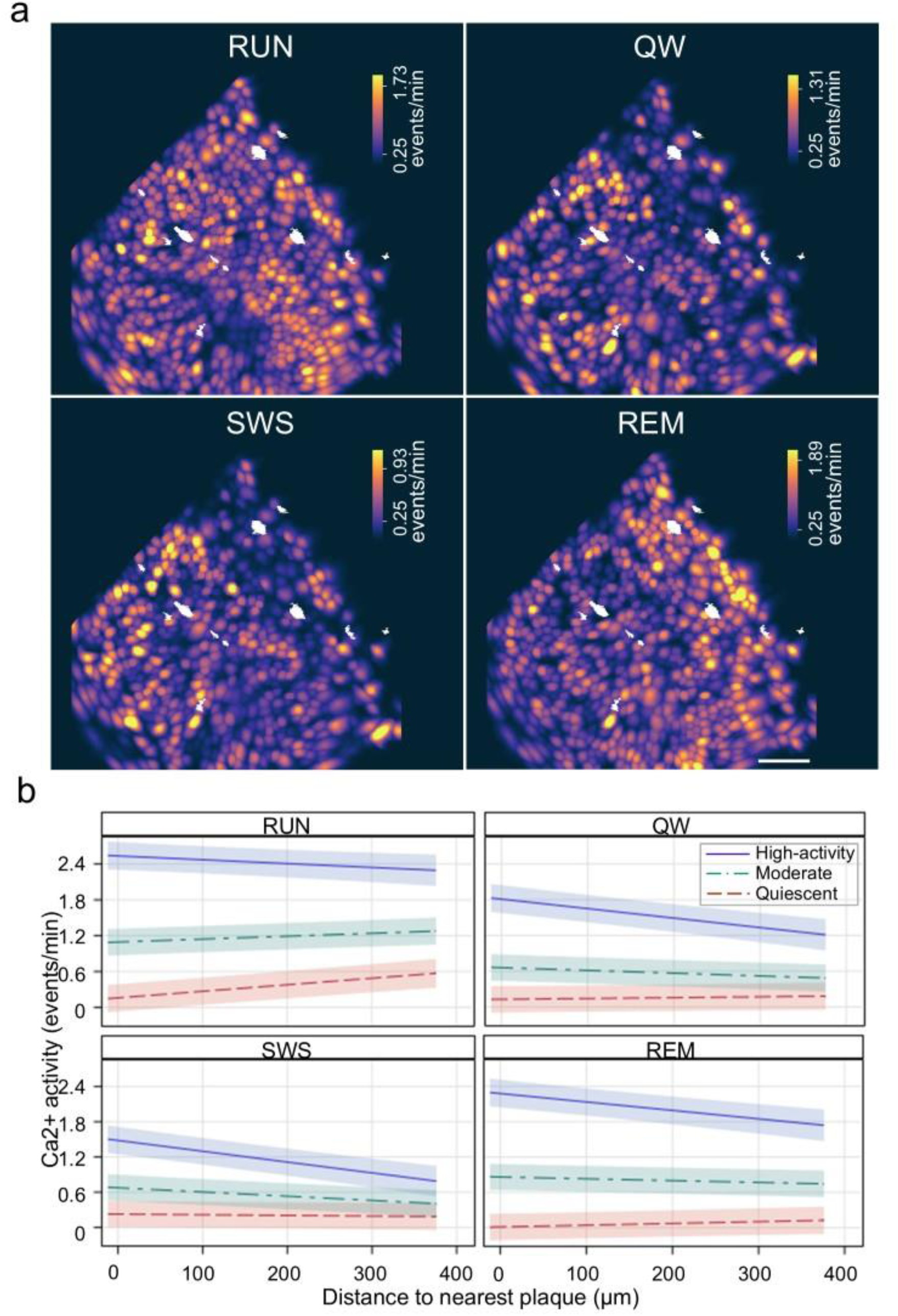
Behavioral state dictates the spatial influence of amyloid plaques on neuronal activity. **a**, Representative spatial maps of hippocampal calcium activity co-registered with amyloid plaques (white) across behavioral states. Color bars indicate the state-specific thresholds for quiescent and high-activity cells. Scale bar, 100 μm. **b,** Variations in calcium activity as a function of distance from the nearest plaque for high-activity, quiescent, and moderately active strata. Proximity to plaques is associated with distinct activity profiles across behavioral states, extending over hundreds of microns (n=7 mice, 3762 cells; mixed effects model, p<0.0001). Data represent the model’s best fit ± 95% confidence intervals.

We next modeled calcium rate as a continuous function of distance to the nearest plaque. Proximity to plaques was associated with higher activity in high-activity cells across states, with effects extending over hundreds of microns, and with lower activity in quiescent cells during running (**Fig. 2b**). Moderately active cells showed a nuanced response, with slightly lower activity near plaques during running and slightly higher activity during QW, SWS, and REM sleep. Importantly, activity class was state-dependent rather than an intrinsic property of the cell, supporting plaque-induced dysfunction as a dynamic circuit-level phenomenon. The specific relationship between plaque proximity and neuronal activity varied markedly across the sleep-wake cycle (**Fig. 2b**). These results suggest that plaques induce differential effects on neuronal activity that depend on both activity type and behavioral state.

## Plaque size and laminar position determine the magnitude of spatial circuit disruption

Amyloid plaques varied in size and laminar position, with some centered in the pyramidal cell layer and others in dendritic layers of stratum radiatum and stratum oriens (**Fig. 3a**). Because amyloid plaques disrupt both the neuronal processes that traverse them^20^ and local inhibitory synapses^21^, we next determined whether plaque size or laminar position within CA1 correlated with specific activity patterns.

**Figure 3.**
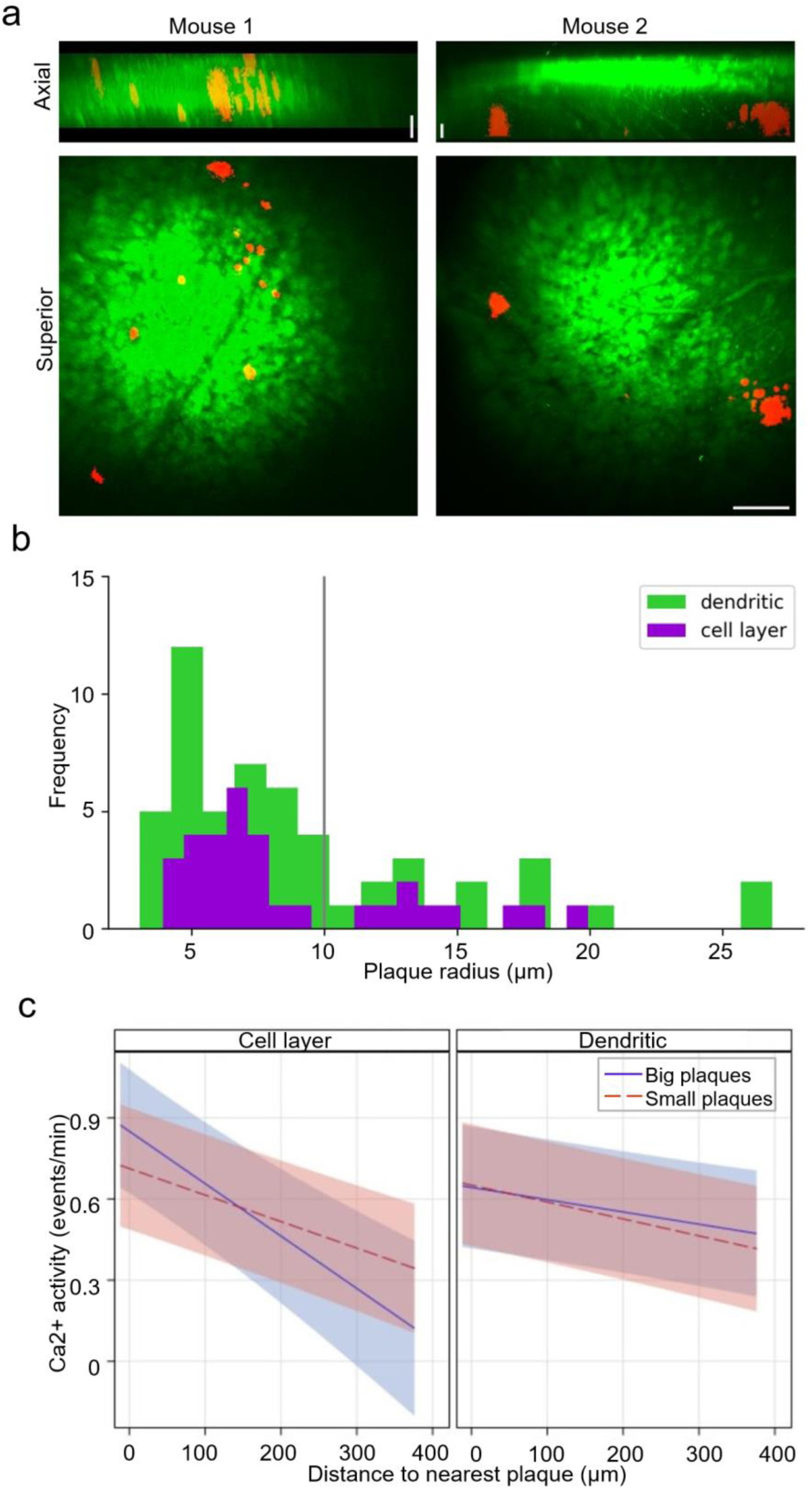
Plaque size and laminar position determine the spatial gradient of neuronal hyperactivity. **a**, Representative 2-photon images of amyloid plaques (red) relative to GCaMP6F-expressing neurons in the CA1 pyramidal cell layer (green). (**Top**) Laminar view distinguishing ‘cell layer’ versus ‘dendritic’ plaques. (**Bottom**) Superior (XY) view. Scale bar, 100µm. **b**, Distribution of plaque diameters for cell layer and dendritic plaque populations (n=7 mice). The solid gray line indicates the threshold determined by k-means clustering. **c**, Spatial decay of calcium activity as a function of plaque diameter and laminar location (n=7 mice, 3762 cells; mixed effects model, p=0.0039). Large plaques in the pyramidal cell layer are associated with the steepest activity gradients. Fits are computed for moderate activity cells during slow-wave sleep (SWS) at a representative edge distance of 136.6 μm. Data represent the model’s best fit ± 95% confidence intervals.

Neuronal activity levels were strongly associated with both plaque size and laminar position (**Fig. 3b**). Large pyramidal-layer plaques were associated with the highest plaque-adjacent activity and the steepest decay with distance. Small pyramidal-layer plaques showed intermediate effects, and dendritic-layer plaques had the smallest impact. Thus, the spatial extent and magnitude of amyloid-driven neuronal dysfunction are shaped by specific plaque characteristics and anatomical location.

## The spatial organization of hippocampal maps relates to amyloid plaque topography

Spatial memory impairment is a hallmark of Alzheimer’s disease^22^ that is well-recapitulated in the APP/PS1 model^3–5^. Given that hippocampal place cell recruitment during navigation is critical for spatial memory formation^23^, we evaluated the impact of amyloid plaques on hippocampal spatial representations. APP/PS1 mice exhibited a lower proportion of place cells than age-matched wild-type controls (**Fig. 4a-c**). Although the percentage of time spent running was similar, APP/PS1 mice ran more slowly, a potential confound that may contribute to reduced place cell yield. Thus, despite reduced place cell yield and altered physiology^3–5,16^, the core hippocampal network required for place coding remained functional in APP/PS1 mice, although future work will be required to disentangle disease effects from speed-related influences on place cell recruitment.

**Figure 4.**
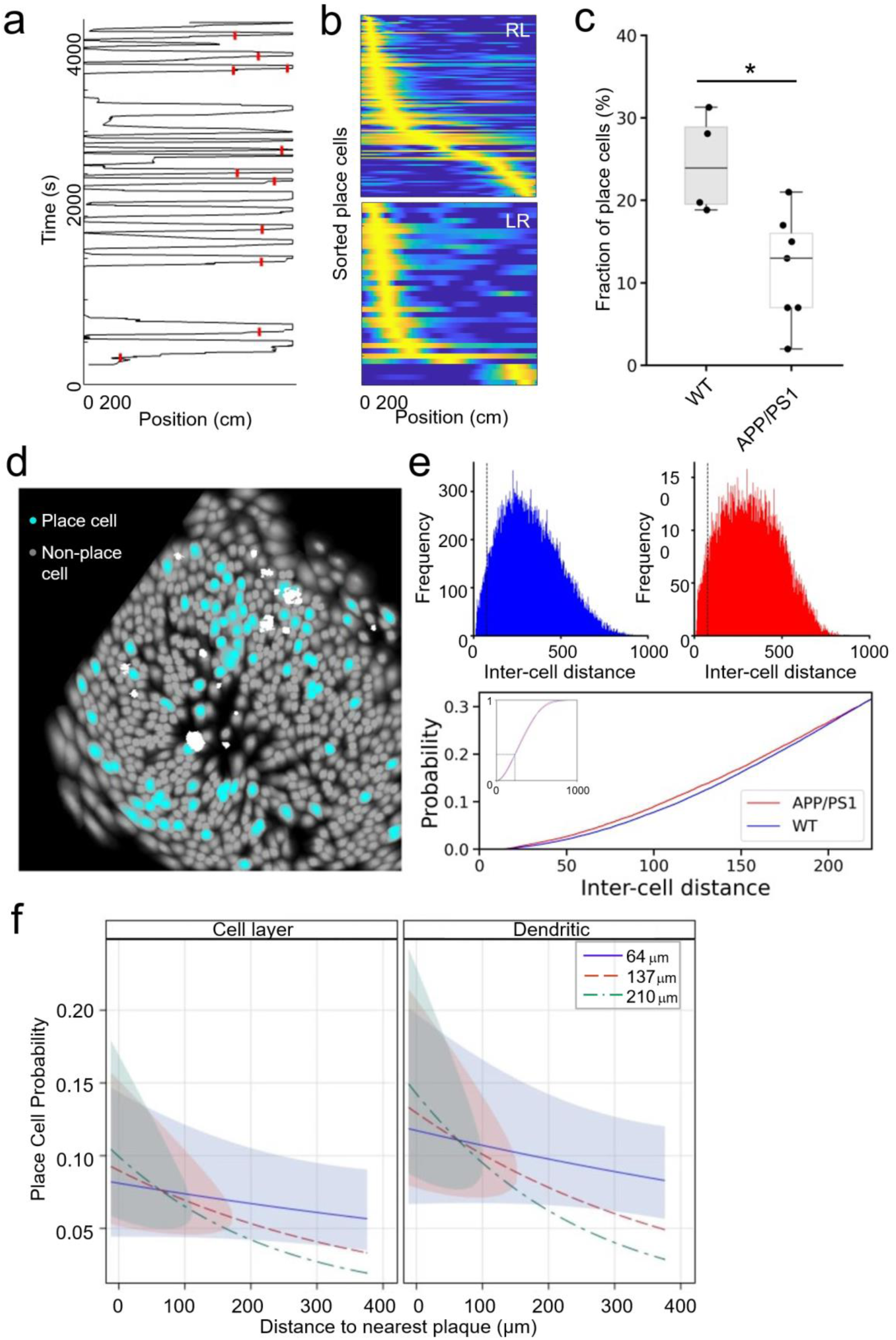
Amyloid plaques are associated with the aberrant spatial clustering of hippocampal place cells. **a**, Spatial tuning of a representative CA1 neuron: Calcium transients (red vertical bars) are plotted against mouse position on a linear track. **b**, Heatmaps of place cell activity sorted by place field location for inbound (left-right; LR) and outbound (right-left; RL) directions. Color bar indicates probability. Yellow indicates the peak field location. **c**, Reduced proportion of place cells in APP/PS1 mice (n=7) compared to age-matched wild-type (WT) controls (n=4). Box plots indicate median (center line), 25^th^/75th percentiles (bounds), and full range (whiskers). **d**, Representative spatial map showing co-registered place cells (cyan), non-place cells (gray), and amyloid plaques (white). **e**, (**Top**) Distribution of inter-cell distances for WT (blue) and APP/PS1 (red) place cells, showing an over-representation of short distances in APP/PS1 mice (vertical line, 75 μm). (**Bottom**) Empirical cumulative distribution function (eCDF) of inter-cell distances (p<10^-14^, Kolmogorov-Smirnov test) (Inset: full eCDF). **f**, In APP/PS1 mice, the probability of a neuron being a place cell increases with proximity to the nearest amyloid plaque (p<0.0001, mixed effects model). This effect is modulated by distance from the edge of the field of view (inset) (mixed effects model, p<0.0001). Place cell probability is higher near dendritic layer plaques than cell layer plaques (p<0.005). * p<0.05. WT: n=4 mice (559 place cells, 2468 total cells); APP/PS1: n=7 mice (410 place cells, 3762 total cells). Data are best fit ± 95% confidence intervals.

To test whether plaques shape place cell topography, we compared the distances between place cells in APP/PS1 and WT mice and related place cell identity to plaque proximity. The distribution of place cell distances in APP/PS1 mice over-represented short distances (**Fig. 4d,e**). In addition, the probability of a neuron being a place cell increased significantly with decreasing distance to the nearest plaque (**Fig. 4f**; see **Methods**). Notably, this effect was greatest near dendritic-layer plaques. In contrast, plaque proximity did not affect place cell spatial information (**Supplementary Fig. 3**), indicating that plaques bias place cell recruitment rather than the quality of their spatial tuning. Together, these results show that place cells are clustered near amyloid plaques.

## Place cell topography does not prefigure future amyloid deposition sites

Neuronal activity drives release of Aβ42^24^ and contributes to its concentration in interstitial fluid^25^. To determine whether place cells aggregate near amyloid plaques or whether pre-existing place cell clusters drive future plaque deposition, we acquired dynamic calcium imaging in CA1 of young APP/PS1 mice prior to plaque formation. We then related the location of these place cells to the topography of future plaques imaged in the same animals 5 months later (**Fig. 5a; see Methods**).

**Figure 5.**
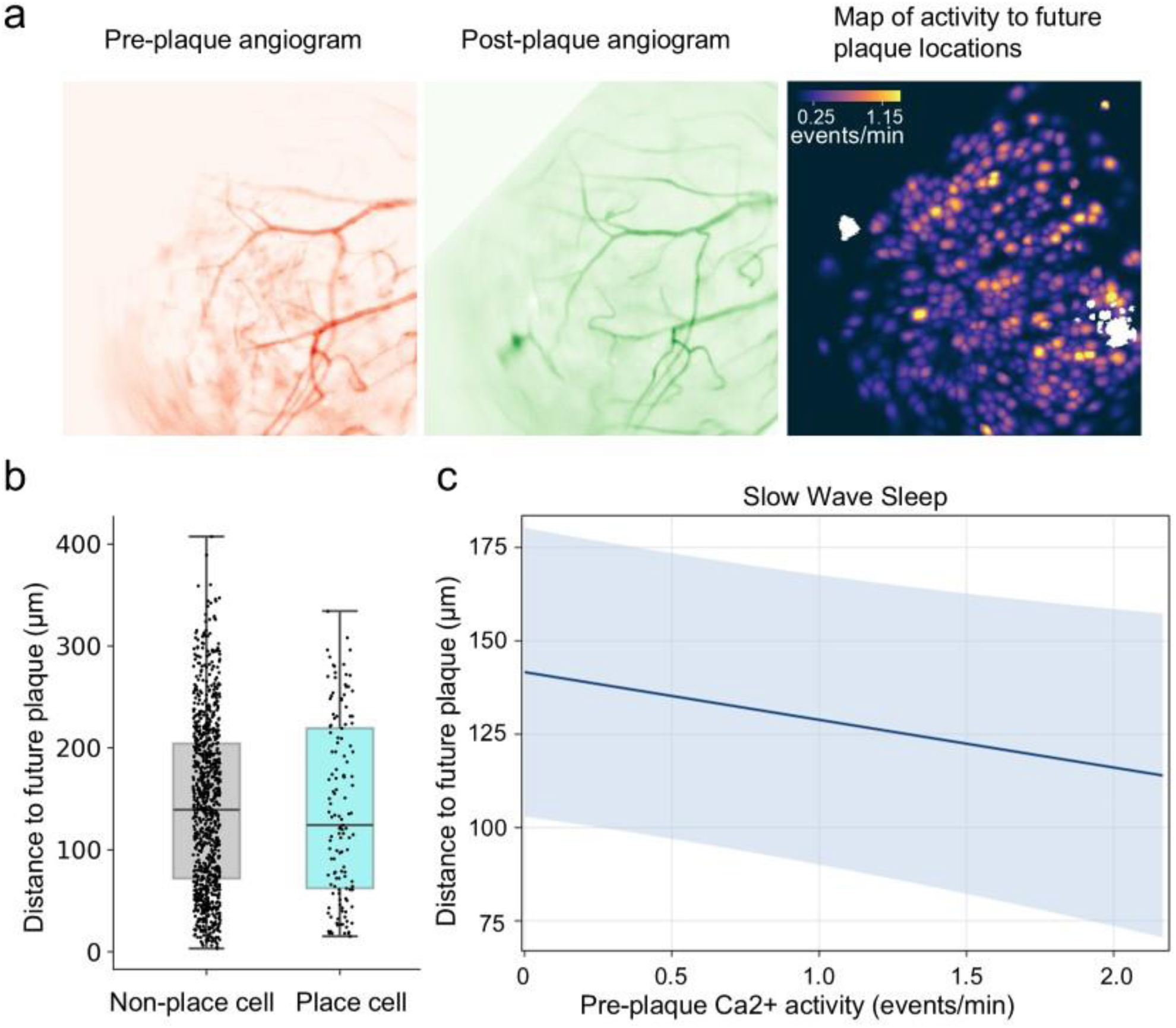
Pre-existing place cell topography does not predict future amyloid deposition sites. **a**, Experimental workflow for longitudinal co-registration. Angiogram fiducials obtained via 1-photon (1P) imaging prior to plaque formation (left) were co-registered with 2-photon (2P) angiograms after plaque deposition (middle) to map initial dynamic calcium activity to future plaque topography. **b**, Proximity to nearest future plaque sites did not differ significantly between place cells and nonplace cells (p>0.05, n.s.). Box plots indicate median (center line), 25^th^/75th percentiles (bounds), and full range (whiskers). **c**, Relationship between pre-plaque neuronal activity and future plaque location. Elevated calcium activity during slow-wave sleep (SWS), but not other behavioral states, was associated with proximity to future amyloid plaques (p=0.038, mixed effects model). Data represent best fit ± 95% confidence intervals (n=3 mice, 1080 cells).

Place cell identity before plaque formation did not predict future plaque locations (**Fig. 5b**). By contrast, elevated neuronal activity during slow-wave sleep was weakly associated with future plaque locations (**Fig. 5c**), suggesting that state-specific hyperactivity during sleep may promote local amyloid deposition. These findings suggest that while the functional architecture of the hippocampal map is reorganized by plaques, its initial layout does not dictate the spatial distribution of amyloid pathology.

## Discussion

Alzheimer’s disease is characterized by progressive hippocampal failure, yet local plaque density correlates poorly with cognitive impairment. Our findings resolve this paradox by showing that amyloid plaques act as drivers of long-range, state-dependent circuit dysfunction in both neuronal activity and the hippocampal map of space. This framework implies that the cognitive burden of the disease reflects network-level collapse due to nonlocal plaque-associated disturbances, rather than simply the volume or density of local plaque deposition.

Place coding is a key function of the hippocampus, and its deterioration in Alzheimer’s patients and in Alzheimer’s models is well established^22^. The biased participation of plaque-adjacent cells in the representation of space reveals a novel mechanism whereby Aβ deposits reorganize existing functional circuits to degrade the functional architecture of the hippocampus. With accumulation of amyloid plaques in Alzheimer’s disease, excessive reliance on a reduced pool of available neurons to represent space within and across environments would be expected to progressively impair both hippocampal remapping and pattern separation, fundamental features of hippocampal mnemonic function^26^.

Our results also suggest that amyloid plaques drive the clustering of place cells rather than vice versa, as our longitudinal studies did not detect a predictive effect of place cell locations on future plaque positions. Although the mechanisms underlying the reorganization of place cells to cluster near amyloid plaques are unknown, loss of inhibitory synapses in the region around amyloid plaques, as recently reported^21^, is well-positioned to contribute, as may the disruption of local processes^20^.

Alternatively, as a smaller proportion of neurons was found to represent space in this Alzheimer’s model compared to WT mice, another possibility is that place coding was lost in cells far from plaques. Unexpectedly, we detected long-range gradients of neuronal activity centered on amyloid plaques rather than highly local plaque effects. Although the biological basis of these effects is not yet clear, and our estimates are necessarily limited by the spatial extent of the imaging window, we anticipate that spatial gradients of plaque-derived soluble Aβ oligomers may contribute, as these species may diffuse through the interstitial space to disrupt synapses far beyond the physical boundaries of the plaque^10,12^.

Prior reports of aberrantly high and low activity of hippocampal and cortical neurons in amyloid plaque-bearing Alzheimer’s models have supported the concept of perturbed excitatory-inhibitory balance in Alzheimer’s disease^1,2,7^. Behavioral state has a marked effect on such imbalance^16^. The present results extend these findings, showing that the spatial effects of plaques on neuronal activity are also markedly state-dependent. In Run behavior, aberrant hyper- and hypoactivity was greatest close to plaques. However, the plaque-associated dysfunction declined faster with distance in non-Run states than in Run, and hypoactivity was unaffected by proximity in non-Run states. Thus, the brain’s functional state dictates the spatial extent of the plaque-associated pathophysiology.

Consistent with the toxicity of Aβ and the local impact of amyloid plaques on the brain parenchyma, large plaques were more disruptive to neuronal activity than small plaques. Interestingly, large plaques deposited in the pyramidal cell layer were most disruptive, while plaques in the dendritic layers were least disruptive. In contrast, we observed that place cells were clustered more around dendritic layer plaques than cell layer plaques. These observations – which suggest a laminar-specific vulnerability of the hippocampal map – may reflect differential effects on inhibitory versus excitatory synapses, given their laminar distribution within CA1^27^ as well as the possible contribution of loss of local inhibitory synapses^21,28^.

Intriguingly, we found that the locations of hyperactive cells in SWS predicted future plaque locations, albeit weakly. As SWS is the one behavioral state associated with amyloid clearance^29,30^, it is possible that release of Aβ in SWS by such hyperactive cells overwhelms local clearance mechanisms, enabling local Aβ concentrations to rise above the local critical threshold for oligomerization and deposition. Future work is needed to evaluate this hypothesis, and larger longitudinal cohorts will be required to determine the robustness of this association.

Key highlights of this study include linking spatial representations and neuronal activity across behavioral states to the local influence of amyloid plaques in the hippocampus, a region critically involved in early Alzheimer’s pathology and memory loss. A further strength was the use of 2P microscopy to image amyloid plaques, providing an accurate assessment of plaque burden across the laminar extent of CA1. While dynamic calcium imaging provides a robust measure of large-scale neuronal ensembles, future studies incorporating high-density electrophysiology may provide additional temporal resolution to the relationship between place cells and amyloid plaques^15,16^. Co-registration of 1P and 2P datasets using angiogram fiducials was associated with low error. Although sample sizes of mice were modest, we were able to sample 85 plaques and 3,762 neurons across behavioral states. Another limitation was the inability to assess the effects of amyloid plaques on calcium in non-somatic cellular compartments. In addition, this work utilized an amyloid-focused APP/PS1 model^31^; future studies in models with robust tau pathology will be important to determine how these circuit-level mechanisms interact with intracellular neurofibrillary pathology.

Together, these findings demonstrate that amyloid plaques are associated with state-dependent, long-range spatial gradients of neuronal activity and a fundamental reorganization of the hippocampal map. Such network-level collapse provides a mechanistic link between focal amyloid deposits and widespread cognitive impairment in Alzheimer’s disease, and suggests general principles by which focal pathology can remap brain circuits in neurodegenerative diseases.

## Methods

All procedures were approved by the Institutional Animal Care and Use Committees of the Massachusetts General Hospital. The study was conducted in accordance with the ethical guidelines of the US National Institutes of Health.

### Surgical preparation

The APP/PS1 mouse strain used for this research project, B6C3-Tg(APPswe,PSEN1dE9)85Dbo/Mmjax, RRID:MMRRC_034829-JAX, was purchased from the Mutant Mouse Resource and Research Center (MMRRC) at The Jackson Laboratory, an NIH-funded strain repository, and was donated to the MMRRC by David Borchelt, Ph.D., McKnight Brain Institute, University of Florida^31^. Surgical preparation has been previously prescribed^32^. Briefly, APP/PS1 mice and their age/sex matched non-transgenic control littermates underwent injection of 1 µL AAV-hSyn-GCaMP6f (Addgene, titer > 7×10^12^ vg/ml) into the hippocampal CA1 region (AP -2.1 mm, ML -1.65 mm, DV 1.4 mm from bregma) under anesthesia (isoflurane 1.5-2%). A gradient index (GRIN) lens (1 mm diameter, AP -2.1 mm, ML -1.50 mm, DV -1.2 mm from bregma) and a custom drivable probe holding 7 stereotrodes (AP -2.8 mm, ML 2.6 mm, -36° relative to the AP axis, -39° relative to the inter-aural axis; DV 1.2 mm) were implanted ipsilaterally 2 weeks later with cerebellar ground. The microscope baseplate was attached three weeks later. Animals were housed in individual cages with a 12 hr light/12hr dark standard light cycle.

### 1P Dynamic calcium imaging and electrophysiology recordings

The Inscopix^TM^ nVista3 mini-microscope and acquisition system (IDAS) were used for dynamic calcium imaging, sampling at 20 fps. Local field potential (LFP) (Neuralynx^TM^) was sampled at 32 kHz followed by 0.5-900 Hz filtering and down sampling to 1000 Hz. Animal position was measured with overhead camera tracking of two LED diodes mounted on the headstage. Electrophysiological recordings and position sampling were acquired continuously. Simultaneous dynamic calcium imaging was acquired in 10-min ON and 5-min OFF blocks to avoid photobleaching and was synchronized with electrophysiological data with TTL pulses.

### Behavior and sleep staging

Data were acquired as animals explored a linear track without reward and were then moved to their home cage within the same recording room to sample subsequent quiet wakefulness (QW), slow-wave sleep (SWS), and REM sleep. Animals were exposed to the track for 1 to 3 days (2.5 ± 0.3 days) before experiments. Cortical delta (1-4 Hz), hippocampal theta (4-10 Hz) and ripples (100-300 Hz), and mouse position and posture were used to identify behavioral states: Run was defined as animals explored the linear track with speeds exceeding 1 cm/sec and was characterized by a strong theta oscillation; QW was identified when animals were stationary (< 0.5 cm/sec) for at least 5 seconds with their eyes open; SWS was identified by the presence of sharp wave ripples (SWRs) and high delta/theta LFP power ratio (>2) in animals in a sleep posture; REM sleep was identified as epochs of high theta/delta LFP power ratio (>2) in animals in a sleep posture^32^.

### 2P imaging of amyloid plaques and acquisition of angiograms

Following 1P imaging of dynamic calcium activity, Methoxy-X04 (10 mg/kg, i.p.) was injected to label amyloid plaques^33^. For 1P vascular imaging at least one day later, animals were then lightly anesthetized with isoflurane (∼1.5%; with concentration sufficient to inhibit dynamic calcium activity), fluorescein dextran (0.3 ml, i.v.) was injected into the tail vein, and imaging was acquired. Animals were then transferred to a 2P microscope and head-fixed for 2P vascular and amyloid plaque imaging on a commercial multiphoton system as previously described^34^ (Olympus Fluoview 1000MPE) mounted on an Olympus BX61WI upright microscope. A mode-locked titanium/sapphire laser (MaiTai; Spectra-Physics, Fremont, CA) was used to generate two-photon fluorescence with 800 nm excitation. Detectors consisting of photomutliplier tubes (Hamamatsu, Ichinocho, Japan) collected light in the following ranges: 380-480 and 500-540 nm. 2P imaging with 800 nm excitation of hippocampal CA1 was acquired through the GRIN lens, acquiring a stack of images across 165 to 344 μm spanning the CA1 cell layer.

### Data analysis

All analyses were performed using MATLAB, Python, or SAS unless otherwise specified^32^.

#### Calcium imaging

Inscopix Data Processing Software (IDPS, Inscopix, Palo Alto, California) was used for movie preprocessing, spatial filtering, motion correction, normalization ΔF/F, and constrained non-negative matrix factorization (CNMFE) cell identification^35^. Each identified cell was manually examined and accepted with oval cell shape and high signal-noise-ratio. Calcium events were then detected at 2 Z scores above the mean, with onset defined at 30% of peak amplitude^32^. Low rate cells were defined as < 0.25 transients/min^32^; high rate cells were defined using a calcium event rate threshold set at the top 5% of calcium rates for each behavioral state of each animal.

#### Place cell detection and analysis

Spatial tuning curves (3 cm bins) were constructed for each cell for each running direction (LR and RL) using all detected calcium events acquired during running (speed > 1 cm/s). Spatial information was measured for each cell as the amount of information (in bits) conveyed per calcium event, and was compared to 1000 shuffled spatial information distributions to test significance (Monte Carlo p-value < 0.05)^36^, and taken as the larger of the LR and the RL trajectories.

#### Single timepoint 2P-1P image co-registration and plaque characterization

Custom Python code was used to co-register 1P and 2P angiograms, implementing a coherent point drift algorithm and Python package pycpd 2.0.0^37,38^. In brief, landmark points marked on 1P and 2P angiograms were used to compute the parameters for a similarity transformation (that included translation, rotation, and scaling) that maximized their overlap. These transformation parameters were then applied to the 1P angiograms and dynamic calcium imaging data to register them in the 2P space. Co-registration accuracy was quantified by calculating the nearest-neighbor distance between corresponding features of the transformed 1P and 2P angiogram datasets. Across the datasets, the root mean square error (RMSE) was 1.9 ± 0.08 µm (mean ± s.e.m.; **Supplementary Fig. 1**). We measured individual plaque size as the cross-sectional area (in μm^2^) from these images using ImageJ software. Plaques were subsequently dichotomized into “small” (radius < 10.7 μm^2^) and “big” (radius ≥ 10.7 μm^2^) categories based on k-means clustering and further dichotomized by their laminar position (pyramidal cell layer vs. dendritic layers; **Fig. 3b**).

#### Young 1P-old 2P image co-registration

The spatial alignment of 1P young calcium imaging data and 2P amyloid plaque image stacks was achieved through a two-stage registration procedure. *Stage 1: Longitudinal 1P image co-registration.* First, a similarity transformation was computed to align the initial angiogram (1P young) with the follow-up angiogram (1P old). Landmark Selection: Corresponding anatomical landmarks were identified on both the 1P young and 1P old angiograms. These landmarks were used to compute the parameters for a first similarity transformation, which included translation, rotation, and uniform scaling. The transformation parameters were optimized to maximize the spatial overlap of the landmark points. The calculated transformation was then applied to the 1P young angiogram and its associated dynamic calcium imaging data, registering them into the coordinate space of the 1P old angiogram. *Stage 2: 2P-1P image co-registration.* To integrate the 1P and 2P datasets, a second registration was performed. A new set of landmarks was selected from major vessels that maintained a consistent structural configuration across the registered 1P young and 2P old angiograms. A second similarity transformation was computed using these stable landmarks to maximize the overlap between the registered 1P young angiogram and the 2P old angiogram. This second set of transformation parameters was then applied to the registered 1P young angiogram and dynamic calcium imaging data to map them into the 2P coordinate space. To evaluate the accuracy of this co-registration method, we computed the nearest-neighbor distance between corresponding features of the transformed 1P young and transformed 1P old angiograms (**Supplementary Fig. 4a**). Across all datasets, the RMSE of this distance was 13.4 ± 0.2 µm (mean ± s.e.m.; **Supplementary Fig. 4d**). The vascular landmarks used for this transformation covered 71.4 ± 3.1% of the total area containing imaged neurons and amyloid plaques (**Supplementary Fig. 4b,c**).

### Statistical analyses of neuronal activity – plaque relationships

#### Neuronal activity as a function of proximity to amyloid plaques

was assessed using a backward eliminated Mixed Effects Model for the dependent variable “Calcium Rate”, using the “State” (Run, QW, SWS, and REM) as a fixed predictor along with Type of Activity (Quiescent, Moderate, and High), the Distance to the nearest plaque, and Edge Distance, including all 2-way interactions. The random effect was Mouse.

The model was significant, revealing several significant 2-way interactions, including of greatest interest, an interaction of distance to the nearest plaque and type of activity (p<0.0001) (**Fig. 2b, Supplementary Fig. 5a**), an interaction of behavioral state and activity type (p<0.0001), an interaction of distance to the nearest plaque and behavioral state (p<0.0001) (**Fig. 2b**), and a shallow but significant interaction of edge distance and activity type (p=0.0065), consistent with the contribution of plaques outside the field of view, such that with greater distance from the edge, the activity of hyperactive cells slightly diminished, the activity of hypoactive cells slightly increased, while the activity of normally active cells remained largely unchanged (**Supplementary Fig. 5b**), indicating that edge distance serves only as a proxy for unobserved plaques beyond the imaging window. The proportion of the variance accounted for by fixed effects in this model was 51.7%.

For **Fig. 1d**, the relationship between distance to the nearest plaque and the proportion of cells in each activity class (high activity, moderate, quiescent) was assessed using Spearman correlations computed separately for each behavioral state. P-values were Bonferroni-adjusted for multiple comparisons.

#### Effect of plaque size and hippocampal laminar position on neuronal activity

was assessed by adding plaque size (big vs. small) and hippocampal laminar position (pyramidal cell layer vs. dendritic layers), their interaction, and the interactions of these two terms with Distance to the nearest plaque and Edge Distance to the previous activity model (which included State, Activity Type, Distance to Plaque, and Edge Distance predictors).

In addition to the same significant 2-way interactions described above, the model detected a significant 3-way interaction of plaque laminar position, plaque size, and distance to the nearest plaque (p=0.0039). The model also revealed a small but significant interaction of plaque laminar position and edge distance (p<0.0001), as well as a modest interaction of activity type and edge distance (p<0.0046) similar to that observed in the previous model (**Supplementary Fig. 6**). The model accounted for 51.8% of the variance.

#### Place cell identity and place cell spatial information as a function of proximity to amyloid plaques

were assessed with two backward elimination mixed effects models: For both, the fixed effect predictors were: Distance to the nearest plaque, the nearest plaque laminar position, nearest plaque size, and Edge Distance (distance to the edge of the field of view, included to account for potential effects of unobserved plaques outside the imaging area), and all two-way interactions. The Random Term was Mouse.

The dependent variable for the first analysis was the binary variable of Place Cell Identity (Yes/No; analyzed via a Generalized Linear Mixed Model assuming a binomial distribution and a logit link function. This analysis revealed a significant interaction between distance to the nearest plaque and edge distance (p<0.0001; **Fig. 4f**). There was also a main effect for plaque laminar position (p=0.005).

The dependent variable for the second analysis was Spatial Information, where we found no significant effect of proximity to amyloid plaques (**Supplementary Fig. 3**).

#### Place cell identity as a function of proximity to future amyloid plaques

was assessed with a mixed effects GLM for the binary dependent variable of pre-plaque Place Cell Identity (Yes/No; measured at timepoint 1). The fixed effect predictor was Distance to Nearest Future Plaque at timepoint 2. The Random Term was Mouse. The GLM assumed a binary distribution and a logit link function for the dependent variable. This analysis showed that place cell identity before plaques formed was not a significant predictor of future plaque location (**Fig. 5b**).

#### Effect of neuronal activity on proximity to future amyloid plaques

was assessed with a mixed effects model for the dependent variable of Distance to Nearest Plaque at timepoint 2. Run, QW, SWS, and REM Calcium Rates (in separate analyses) were used as fixed predictors. The Random Term was Mouse. This analysis showed that higher activity in SWS – but not in Run, QW, or REM sleep – was weakly predictive of future plaque locations (p=0.038; **Fig. 5c**).

## Author contributions

Conceptualization, Z.Z. and S.N.G.

Methodology, Z.Z., R.J.G, K.K., B.B., and S.N.G.

Software, Z.Z., H.L. N.G., and S.N.G.

Validation, Z.Z., H.L., and J.J.L.

Formal Analysis, Z.Z., J.J.L., H.L.

Investigation, Z.Z., H.L., and S.N.G.

Resources, Z.Z., K.K., B.B., and S.N.G.

Data Curation, Z.Z., H.L., and J.J.L. Writing – Original Draft, Z.Z., and S.N.G.

Writing – Review & Editing, Z.Z., H.L., J.L., N.G., R.J.G., K.K., B.B., and S.N.G.

Visualization, Z.Z., H.L., J.J.L.

Supervision, S.N.G.

Project Administration, S.N.G.

Funding Acquisition, S.N.G.

## Data and materials availability

The datasets generated and analyzed in the current study will be available in the Harvard Dataverse repository.

## Extended Data Figures

**Supplementary Figure 1.**
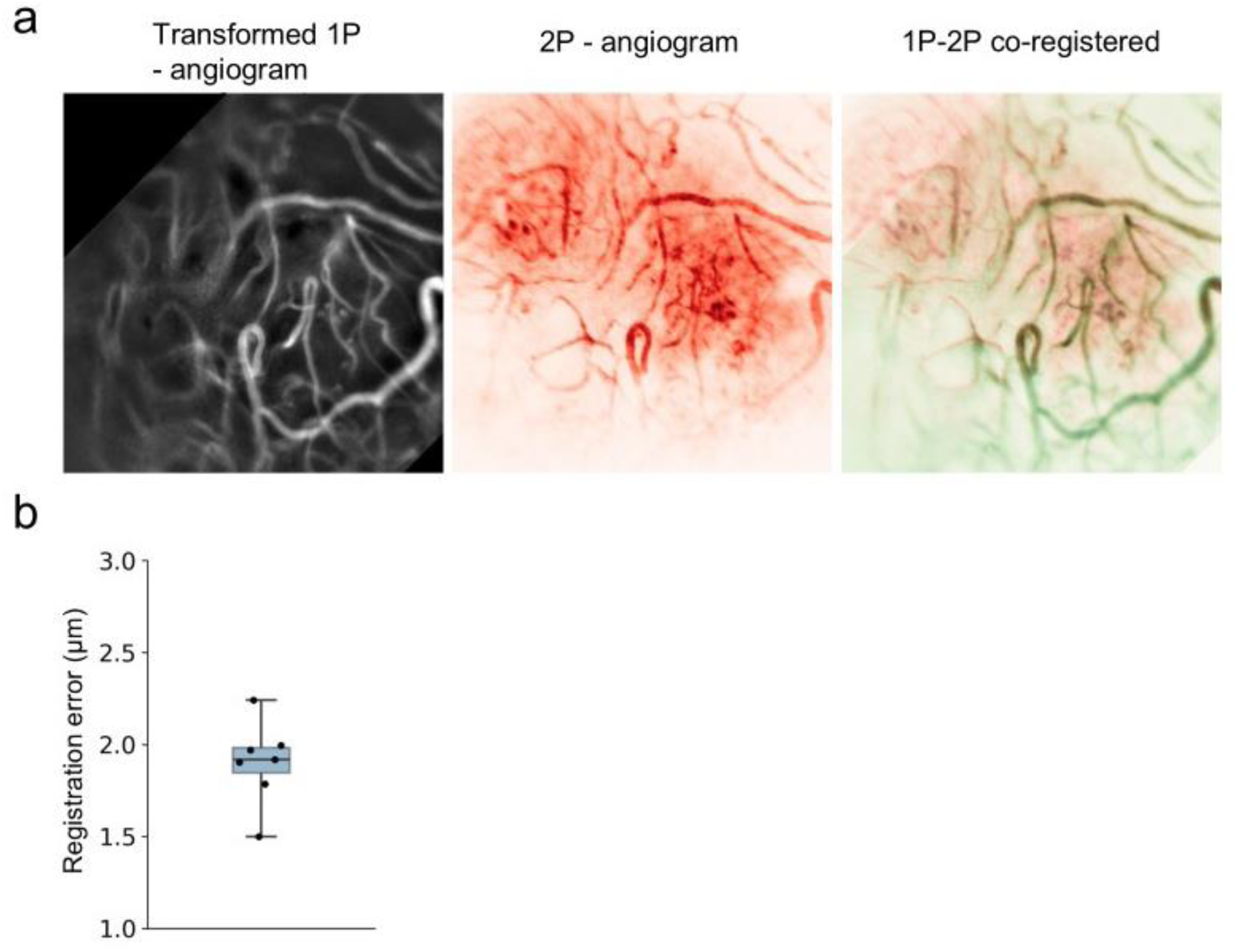
Spatial co-registration of 1-photon and 2-photon imaging volumes. **a**, **Representative co-registration of 1-photon (**1P) neuronal maps and 2-photon (2P) plaque topography. Volumes were aligned via rigid transformation using hippocampal angiogram fiducials. **b**, Precision of co-registration as measured by the root mean square error (RMSE) of the distances between transformed 1P and imaged 2P vasculature (n = 7 co-registrations). Box plot indicate median (center line), 25^th^/75th percentiles (bounds), and full range (whiskers).

**Supplementary Figure 2.**
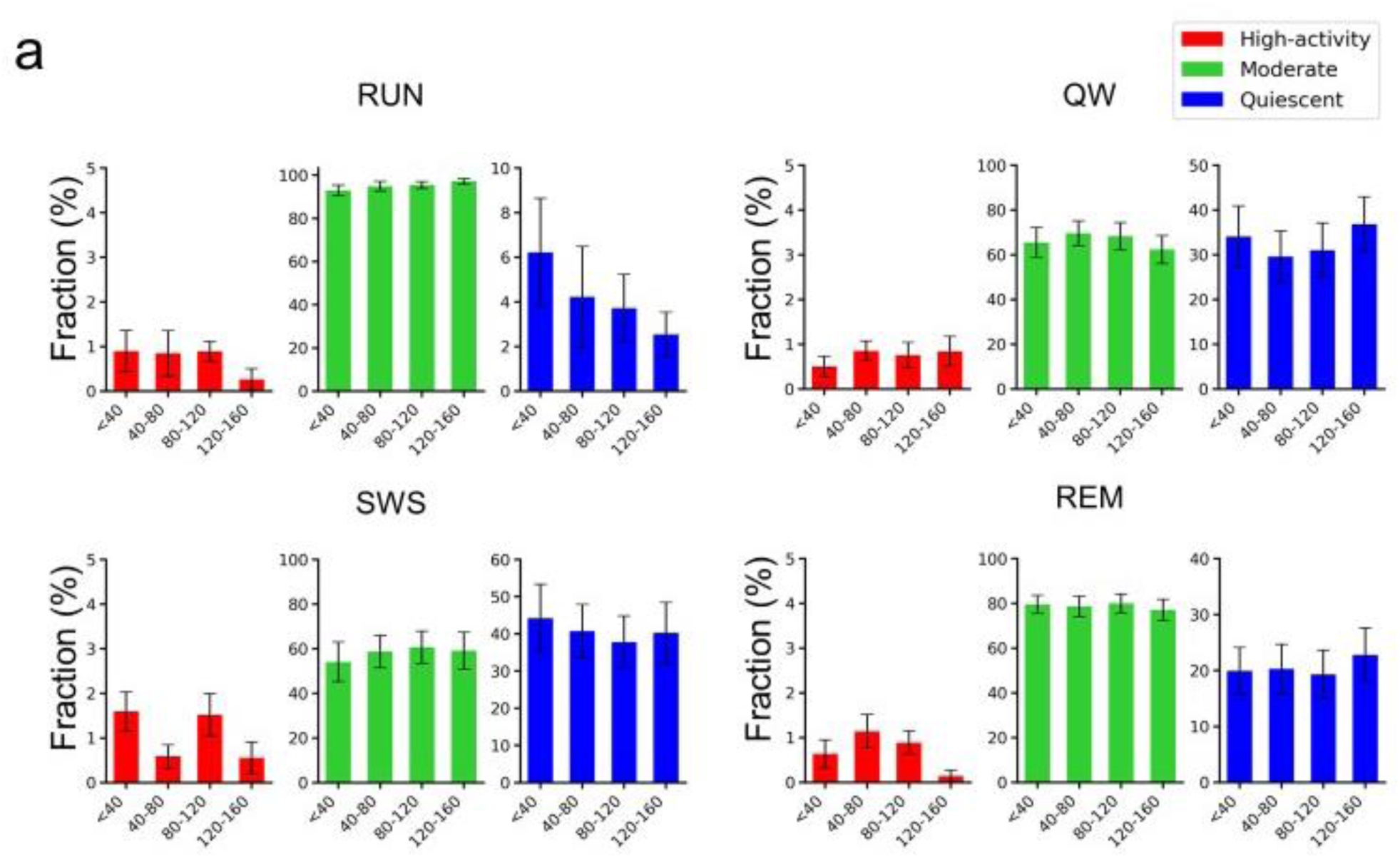
Distribution of activity strata relative to plaque proximity using a 1% hyperactivity threshold. Proportion of high-activity (red), moderate (green), and quiescent (blue) neurons as a function of distance from the nearest plaque across behavioral states. In this analysis, high-activity cells are defined for each state as those cells in the top 1% of calcium event rates. Data are presented as mean ± s.e.m. (n=7 mice, 3,762 cells).

**Supplementary Figure 3.**
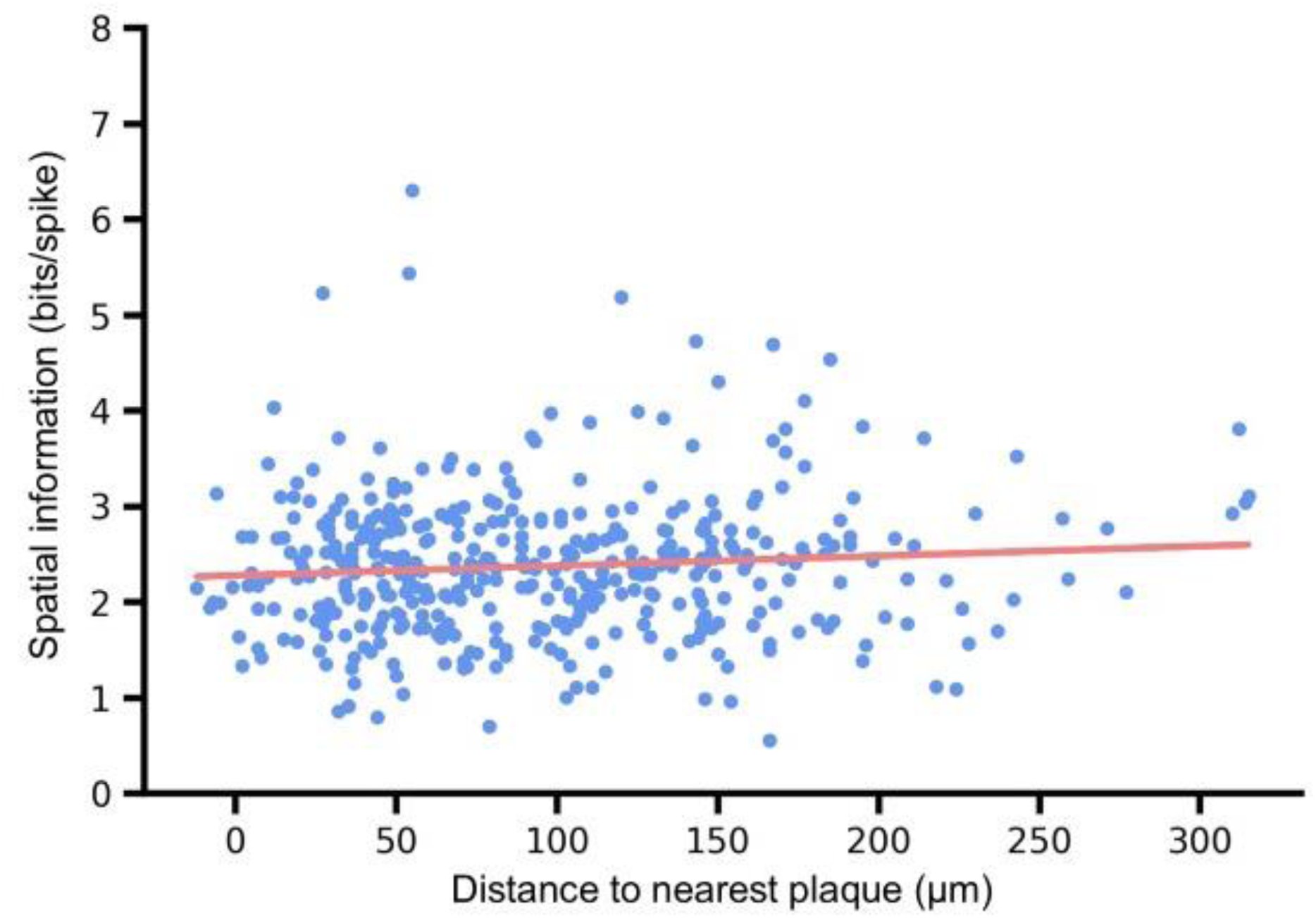
Place cell spatial information does not depend on proximity to amyloid plaques. Linear least squares regression (r = 0.086, p=0.08; red line) and linear mixed-effects modeling (p>0.05) reveal no relationship between a neuron’s spatial information content and its distance to the nearest plaque. Data represent individual place cells (n=410 cells from 7 mice).

**Supplementary Figure 4.**
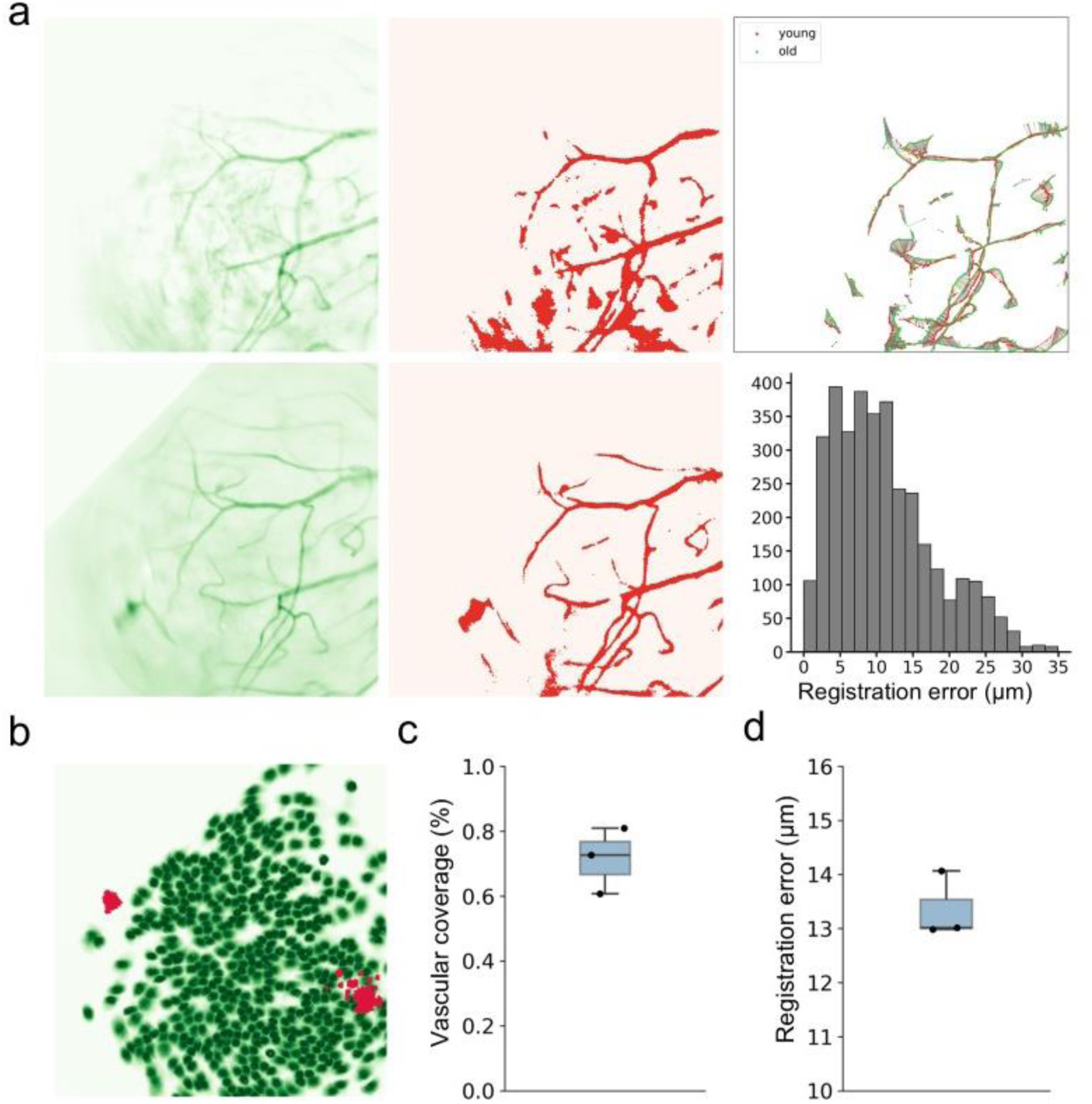
Longitudinal co-registration of hippocampal activity and future plaque topography. **a**, Mapping of hippocampal vasculature across time-points to align pre-plaque calcium activity with future plaque locations. Left: Representative 1-photon (1P) angiograms from a single mouse at 6.2 months (top) and 10.5 months of age (bottom). Middle: Automated identification of fiducial vessels. (Right) Superimposition of identified vessels using rigid body transformation (top) and the resulting spatial registration error (bottom). **b**, Representative field showing co-registered neuronal somas (6.2 months) and amyloid plaques (10.5 months). **c**, Vascular coverage across the co-registered field of view, illustrating the extent of the common imaging area. **d**, Precision of the longitudinal alignment as measured by the root mean square error (RMSE) of the distances between transformed young and old 1P vessels. Box plots indicate median (center line), 25^th^/75th percentiles (bounds), and full range (whiskers).

**Supplementary Figure 5.**
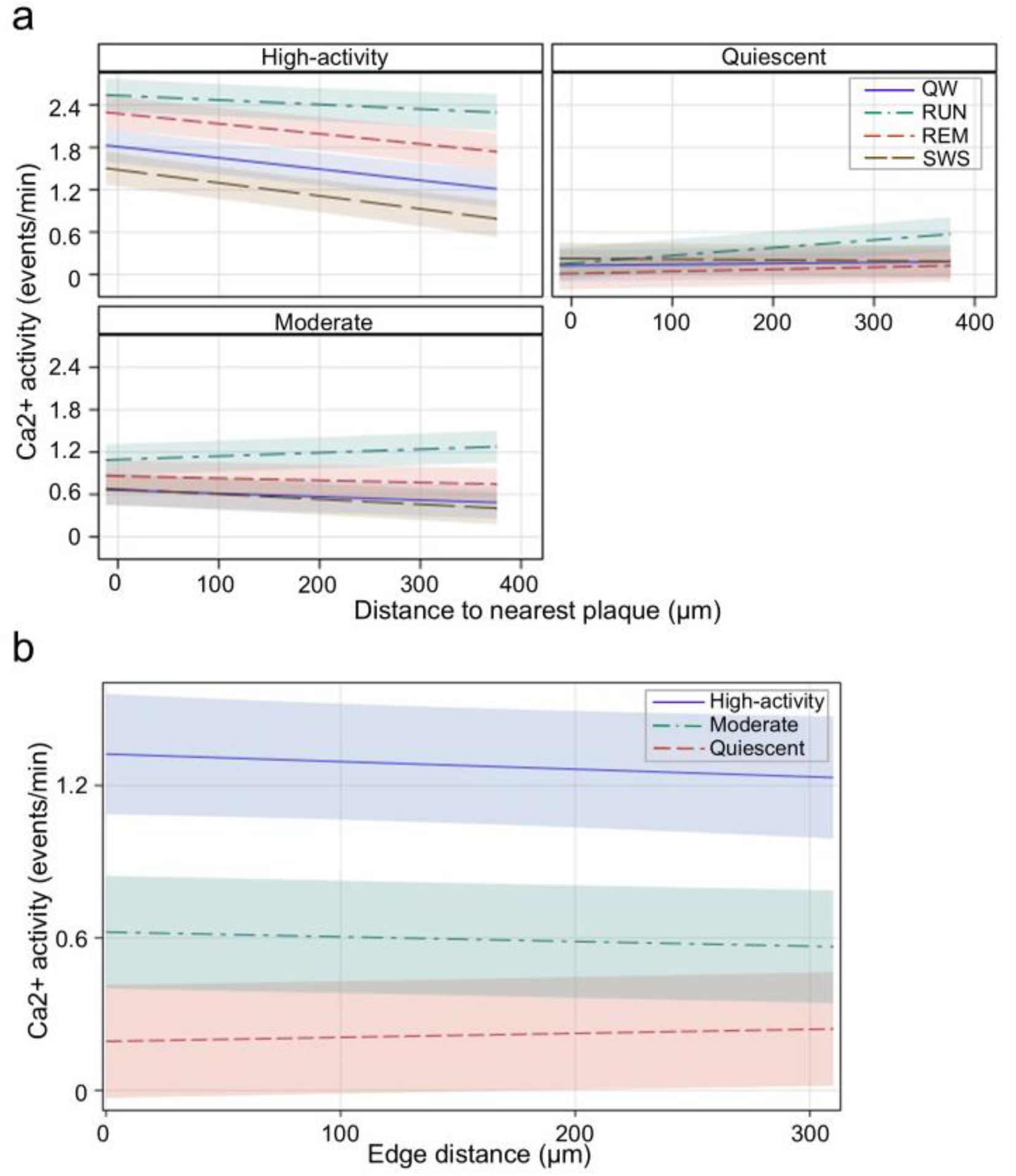
State-dependent spatial gradients and edge-effect modeling of calcium activity. **a**, High activity, moderate, and quiescent neurons exhibit distinct activity profiles as a function of distance from the nearest plaque across behavioral states ( n=7 mice, 3,762 cells; linear mixed effects model, p<0.0001). **b**, Calcium activity as a function of distance from the Edge of the field of view for each activity stratum (linear mixed effects model, p=0.0065). Data represent the model’s best fit ± 95% confidence intervals.

**Supplementary Figure 6.**
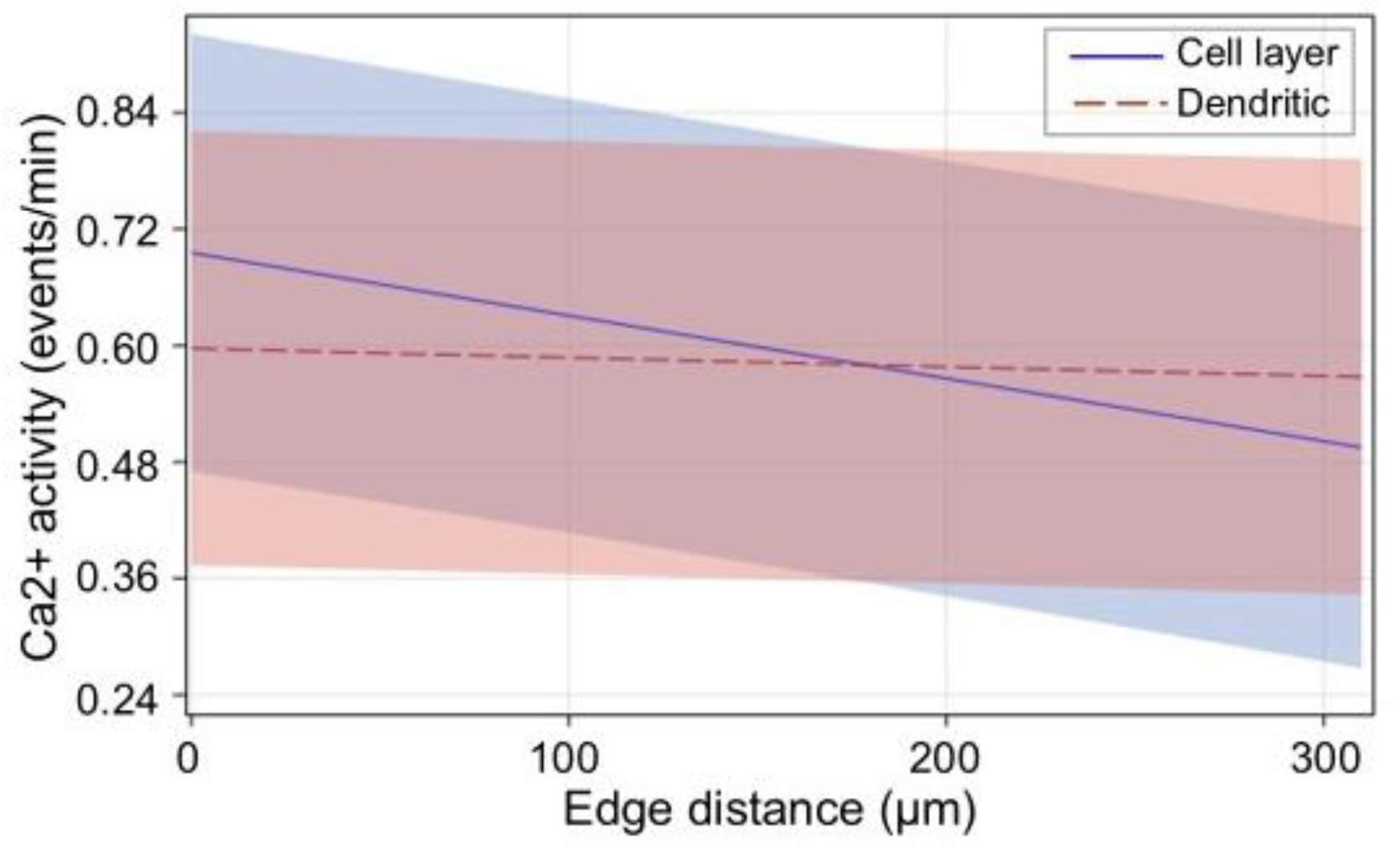
Interaction between plaque laminar position and field-of-view edge distance. Calcium activity as a function of distance from the Edge of the field of view, stratified by the laminar position (cell layer vs. dendritic) of the nearest amyloid plaque (p<0.0001, linear mixed effects model). Data represent the model’s best fit ± 95% confidence intervals.

